# Smaller spared subcortical nuclei are associated with worse post-stroke sensorimotor outcomes in 28 cohorts worldwide

**DOI:** 10.1101/2020.11.04.366856

**Authors:** Sook-Lei Liew, Artemis Zavaliangos-Petropulu, Nicolas Schweighofer, Neda Jahanshad, Catherine E. Lang, Keith R. Lohse, Nerisa Banaj, Giuseppe Barisano, Lee A. Baugh, Anup K. Bhattacharya, Bavrina Bigjahan, Michael R. Borich, Lara A. Boyd, Amy Brodtmann, Cathrin M. Buetefisch, Winston D. Byblow, Jessica M. Cassidy, Valentina Ciullo, Adriana B. Conforto, Richard C. Craddock, Adrienne N. Dula, Natalia Egorova, Wuwei Feng, Kelene A. Fercho, Chris M. Gregory, Colleen A. Hanlon, Kathryn S. Hayward, Jess A. Holguin, Brenton Hordacre, Darryl H. Hwang, Steven A. Kautz, Mohamed Salah Khlif, Bokkyu Kim, Hosung Kim, Amy Kuceyeski, Bethany Lo, Jingchun Liu, David Lin, Martin Lotze, Bradley J. MacIntosh, John L. Margetis, Feroze B. Mohamed, Jan Egil Nordvik, Matthew A. Petoe, Fabrizio Piras, Sharmila Raju, Ander Ramos-Murguialday, Kate P. Revill, Pamela Roberts, Andrew D. Robertson, Heidi M. Schambra, Na Jin Seo, Mark S. Shiroishi, Surjo R. Soekadar, Gianfranco Spalletta, Cathy M. Stinear, Anisha Suri, Wai Kwong Tang, Gregory T. Thielman, Vincent N. Thijs, Daniela Vecchio, Junping Wang, Nick S. Ward, Lars T. Westlye, Carolee J. Winstein, George F. Wittenberg, Kristin A. Wong, Chunshui Yu, Steven L. Wolf, Steven C. Cramer, Paul M. Thompson, on behalf of the ENIGMA Stroke Recovery Working Group

## Abstract

**Background and Purpose:** Up to two-thirds of stroke survivors experience persistent sensorimotor impairments. Recovery relies on the integrity of spared brain areas to compensate for damaged tissue. Subcortical regions play critical roles in the control and regulation of sensorimotor circuits. The goal of this work is to identify associations between volumes of spared subcortical nuclei and sensorimotor behavior at different timepoints after stroke.

**Methods:** We pooled high-resolution T1-weighted MRI brain scans and behavioral data in 828 individuals with unilateral stroke from 28 cohorts worldwide. Cross-sectional analyses using linear mixed-effects models related post-stroke sensorimotor behavior to non-lesioned subcortical volumes (Bonferroni-corrected, p<0.004). We tested subacute (≤90 days) and chronic (≥180 days) stroke subgroups separately, with exploratory analyses in early stroke (≤21 days) and across all time. Sub-analyses in chronic stroke were also performed based on class of sensorimotor deficits (impairment, activity limitations) and side of lesioned hemisphere.

**Results:** Worse sensorimotor behavior was associated with a smaller ipsilesional thalamic volume in both early (n=179; *d*=0.68) and subacute (n=274, *d*=0.46) stroke. In chronic stroke (n=404), worse sensorimotor behavior was associated with smaller ipsilesional putamen (*d*=0.52) and nucleus accumbens (*d*=0.39) volumes, and a larger ipsilesional lateral ventricle (*d*=-0.42). Worse chronic sensorimotor impairment specifically (measured by the Fugl-Meyer Assessment; n=256) was associated with smaller ipsilesional putamen (*d*=0.72) and larger lateral ventricle (*d*=-0.41) volumes, while several measures of activity limitations (n=116) showed no significant relationships. In the full cohort across all time (n=828), sensorimotor behavior was associated with the volumes of the ipsilesional nucleus accumbens (*d*=0.23), putamen (*d*=0.33), thalamus (*d*=0.33), and lateral ventricle (*d*=-0.23).

**Conclusions:** We demonstrate significant relationships between post-stroke sensorimotor behavior and reduced volumes of subcortical gray matter structures that were spared by stroke, which differ by time and class of sensorimotor measure. These findings may provide additional targets for improving post-stroke sensorimotor outcomes.

## INTRODUCTION

Sensorimotor recovery after stroke relies on residual motor architecture.^1^ The majority of research in this area has focused on the role of cortical regions within sensorimotor networks, which often undergo significant reorganization and vary widely across individuals after stroke. Spared subcortical nuclei also form key components of corticothalamic and corticostriatal circuits that support sensorimotor performance but have been less studied in recent years. These structures may yield additional insight into processes impacting stroke outcomes, given their clearly defined boundaries, well-mapped inputs and outputs, and known associations with specific neurotransmitters and genetic variants.^2^

As relay nodes for sensorimotor circuits in the brain, subcortical nuclei not only play a critical role in the maintenance and regulation of networks for motor learning, but they also subserve cognition, metabolic regulation, and reward—all of which have been implicated as contributors to post-stroke outcomes, including sensorimotor functioning and recovery.^3-6^ Each structure in the cortico-striatal-thalamic circuit has a distinct role in sensorimotor control and possibly outcomes. For instance, the thalamus is integral to the regulation of metabolism, sleep and wakefulness, cognitive processing, and integrating sensorimotor information,^7^ and thalamic metabolism has been shown to be disordered in the early weeks after stroke.^3, 8^ Similarly, the basal ganglia (e.g., caudate, putamen, globus pallidus, and nucleus accumbens) are heavily involved in motor control, learning, and reward, with distinct roles for each nuclei.^9, 10^ Direct damage to the thalamus and basal ganglia is associated with poor sensorimotor behavior and recovery,^4, 11^ but the role of each spared subcortical nuclei is unclear.

To date, these subcortical structures have been studied only in modestly-sized samples, with varying results, and with measurements from multiple regions often aggregated as one (e.g., combined analysis of the thalamus and basal ganglia). However, each nucleus has a characteristic distribution of neurotransmitters and network connections; identifying specific non-lesioned subcortical nuclei could provide more precise neurobiological targets for therapeutics to potentiate recovery.

In addition, inter-individual variability and the heterogeneity of brain changes after stroke pose challenges to the identification of neural targets in spared tissue. Addressing this issue requires large, diverse, and appropriately powered sample sizes with high-resolution brain MRIs. Although acute stroke research has successfully utilized pooled approaches with individual patient data to examine acute treatment outcomes,^12, 13^ stroke rehabilitation research has been slower to adopt this type of approach due to the complexity of combining elaborate rehabilitation research protocols, differences in the site and size of infarcts, diversity of the patient populations recruited, and variety of the stroke neuroimaging and behavioral measures collected. To address these challenges, we formed the international ENIGMA Stroke Recovery Working Group to harmonize and combine diverse individual patient data, including high-resolution structural brain MRIs and behavioral outcome measures, across multiple research centers.^14^ This combined analysis pools individual patient data across research sites using a harmonized analytical pipeline and includes both published and unpublished data. Compared to traditional single-site analyses or retrospective meta-analyses, this approach allows for greater statistical rigor, testing of more sophisticated hypotheses (e.g., subgroup analyses), and less bias due to the inclusion of both published and unpublished data across diverse cohorts.^15, 16^ Furthermore, pooled analyses with multi-site data increase heterogeneity, which improves generalizability of findings, reduces research inefficiency by leveraging previously collected data to examine novel questions, and advances the field faster than is achievable by prospective studies.^17^

The current study pools data from 828 individuals across 28 cohorts worldwide from the ENIGMA Stroke Recovery Working Group to examine relationships between sensorimotor behavioral measures and volumes of the ipsilesional and contralesional thalamus, putamen, caudate, pallidum, and nucleus accumbens. Enlargement of the lateral ventricles was also examined as an indirect measure of atrophy and vascular integrity.^18, 19^ Given the neurobiological events unique to early and subacute stroke compared to chronic stroke, data were analyzed separately for individuals in the subacute (≤ 90 days) and chronic (≥ 180 days) stages.^20^ As an exploratory measure, we also analyzed relationships early after stroke (≤ 21 days), before post-stroke secondary structural atrophy is thought to be observed,^21^ to estimate whether subacute associations are driven by early post-stroke changes or likely existed prior to the stroke, as well as across all time.

We hypothesized that thalamic volume would relate to sensorimotor behavior in early and subacute phases after stroke, given its multiple roles in supporting cellular repair.^3, 22^ We further expected that smaller subcortical volumes (reflecting atrophy of structures associated with sensorimotor control) and larger ventricles (reflecting general atrophy) would be related to worse chronic sensorimotor behavior.^23^

Furthermore, as sensorimotor behavior encompasses multiple classes of the International Classification of Functioning, Disability, and Health (ICF), we conducted separate subgroup analyses in chronic stroke to examine if there are specific neural correlates of loss of body structures and function (i.e., *sensorimotor impairment*) versus loss of activity in daily tasks (i.e., *activity limitations*).^24^ We anticipated that subcortical nuclei important for direct sensorimotor control, such as the putamen, would more strongly relate to impairment; conversely, regions associated with reward and motivation, such as the nucleus accumbens, should more strongly relate to activity limitation. Finally, in chronic stroke, we also examined the impact of the side of the lesion. Based on evidence of hemispheric specialization for motor behavior after stroke,^25^ we hypothesized that the side of the lesion would modify the relationship between non-lesioned subcortical tissue volume and sensorimotor behavior.

## MATERIALS AND METHODS

### Study design

The current cross-sectional pooled analysis used data from the ENIGMA Stroke Recovery Working Group, which was frozen for this analysis on May 22, 2020. A detailed overview of ENIGMA Stroke Recovery procedures and methods are reported elsewhere.^14^ The retrospective data were collected across 28 different research studies (i.e., cohorts) at 16 different research institutes in 10 countries. Data were collected in accordance with the Declaration of Helsinki and in compliance with local ethics review boards at each institute (see *Supplementary Table 1* for details).

**Table 1.** Summary of research cohort characteristics. Age and sensorimotor behavioral score data are shown as median (interquartile range (IQR), minimum-maximum values)

| **Cohort ID** | ***n*** | **Females / Males** | **Median Age**<br>**(IQR, min-max)** | **Median Sensorimotor Score**<br>**(IQR, min-max)** |
| --- | --- | --- | --- | --- |
| 1 | 39 | 10 / 29 | 61 (17, 31-80) | 0.65 (0.23, 0.0-0.9) |
| 2 | 12 | 06 / 06 | 70 (12, 39-85) | 0.50 (0.41, 0.2-0.7) |
| 3 | 14 | 06 / 08 | 60 (15, 33-85) | 0.25 (0.22, 0.1-0.6) |
| 4 | 19 | 06 / 13 | 44 (15, 30-68) | 0.14 (0.17, 0.0-0.5) |
| 7 | 42 | 14 / 28 | 56 (14, 18-80) | 0.82 (0.35, 0.4-1.0) |
| 8 | 8 | 02 / 06 | 62 (10, 39-75) | 0.55 (0.35, 0.0-1.0) |
| 9 | 93 | 29 / 64 | 70 (16, 24-88) | 1.00 (0.07, 0.0-1.0) |
| 10 | 24 | 05 / 19 | 59 (13, 42-74) | 1.00 (0.02, 0.7-1.0) |
| 11 | 29 | 10 / 19 | 57 (11, 44-71) | 1.00 (0.05, 0.1-1.0) |
| 12 | 57 | 31 / 26 | 71 (17, 31-97) | 0.65 (0.71, 0.0-1.0) |
| 13 | 44 | 22 / 22 | 72 (18, 33-91) | 0.12 (0.32, 0.0-1.0) |
| 15 | 14 | 06 / 08 | 57 (11, 45-74) | 0.72 (0.25, 0.4-0.8) |
| 17 | 16 | 05 / 11 | 59 (04, 45-68) | 0.55 (0.23, 0.2-0.7) |
| 18 | 11 | 05 / 06 | 59 (07, 46-73) | 0.65 (0.22, 0.5-0.9) |
| 19 | 13 | 03 / 10 | 62 (21, 33-74) | 0.84 (0.08, 0.8-0.9) |
| 20 | 22 | 08 / 14 | 70 (13, 49-79) | 0.91 (0.14, 0.3-1.0) |
| 22 | 17 | 04 / 13 | 59 (30, 25-72) | 0.63 (0.50, 0.0-0.8) |
| 23 | 13 | 07 / 06 | 58 (08, 31-90) | 0.42 (0.17, 0.3-0.8) |
| 24 | 21 | 11 / 10 | 63 (13, 32-78) | 0.95 (0.00, 0.6-1.0) |
| 25 | 26 | 10 / 16 | 65 (18, 37-88) | 0.97 (0.20, 0.0-1.0) |
| 26 | 24 | 14 / 10 | 49 (20, 25-71) | 0.64 (0.14, 0.3-0.8) |
| 28 | 26 | 07 / 19 | 62 (11, 23-75) | 0.75 (0.25, 0.3-1.0) |
| 31 | 35 | 09 / 26 | 58 (12, 21-86) | 0.52 (0.31, 0.2-0.9) |
| 32 | 7 | 03 / 04 | 62 (16, 38-72) | 0.95 (0.44, 0.2-1.0) |
| 34 | 15 | 06 / 09 | 58 (11, 32-80) | 0.82 (0.20, 0.6-1.0) |
| 35 | 15 | 06 / 09 | 64 (18, 31-83) | 0.64 (0.52, 0.2-0.9) |
| 38 | 81 | 34 / 47 | 66 (19, 30-89) | 0.85 (0.60, 0.0-1.0) |
| 41 | 91 | 33 / 58 | 70 (15, 32-89) | 1.00 (0.02, 0.8-1.0) |
| **TOTAL** | **828** | **312 / 516** | **63 (19, 18-97)** | **0.82 (0.48, 0-1)** |

### ENIGMA Stroke Recovery Dataset

Participants with at least one sensorimotor behavioral outcome measure (see *Behavioral Data Analysis*) and a segmented high-resolution (e.g., 1-mm isotropic) T1-weighted (T1w) structural MRI of the brain (see *MRI Data Analysis*) were included, yielding an initial dataset of 1,285 individuals. Only participants with unilateral ischemic stroke or intracerebral hemorrhage were included, and individuals identified as having bilateral lesions or lesions in the brainstem or cerebellum were excluded from this analysis. For any longitudinal observations, only the first time-point was used; the resulting dataset was therefore cross-sectional. Each brain region was manually inspected for quality and overlap with the lesion (see *MRI Data Analysis)*. Any individuals missing covariates of age (n=50) or sex (n=89) were also excluded, yielding a final sample of 828 individuals. As the relationships between brain volume and sensorimotor behavior were expected to change with time after stroke, the data were divided into subacute stroke (≤90 days post-stroke) and chronic stroke (≥180 days post-stroke). Exploratory analyses looking only at early stroke (≤21 days post-stroke) and across all times after stroke are also included.

### MRI Data Analysis

To extract subcortical volumes, the brain imaging software package FreeSurfer (version 5.3) was used to segment subcortical regions of interest (ROIs) from the T1w MRIs.^26^ Twelve ROIs were extracted: the left and right thalamus, caudate, putamen, pallidum, nucleus accumbens, and lateral ventricles. For all analyses, these were characterized as ipsilesional and contralesional with respect to the lesioned hemisphere. Total intracranial volume (ICV) was also quantified using FreeSurfer outputs. ENIGMA scripts developed in-house were used to extract the volume of each ROI for each individual and to generate quality control (QC) triplanar images of each segmented ROI as done previously (http://enigma.ini.usc.edu/protocols/).^2^ Given the variability of post-stroke neuroanatomy following a lesion, trained research team members (A.Z.-P., A.S.) performed visual QC for each ROI in each subject. Any regions intersecting the lesion were marked “lesioned,” and any regions not properly segmented by FreeSurfer were marked “failed.” Regions falling in either category were excluded from further analysis (for the full QC protocol, see Appendix 1 in ref^14^). Sample sizes for each analysis and brain region are reported in each results table.

### Behavioral Data Analysis

Across cohorts, behavioral data were collected within approximately 72 hours of the MRI. To maximize the utility of the full dataset, a *primary sensorimotor behavior score* was defined for each study cohort using the measure reported in that cohort that was most commonly represented in the dataset overall (see *Supplementary Materials*). From this measure, a fraction of the maximum possible score was calculated, such that 0 represented the worst sensorimotor performance (severe deficits) and 1 represented the best sensorimotor performance (no deficits). The most common measure across cohorts was the Fugl-Meyer Motor Assessment of Upper Extremities (FMA-UE).^27^

In chronic stroke, we also examined behavioral measures that specifically captured impairment and activity limitation. Impairment was measured by the FMA-UE, whereas activity limitation was measured by the Action Research Arm Test (ARAT)^28^ and Wolf Motor Function Test (WMFT).^29^ These data were not examined in early stroke due to the limited sample sizes with these measures.

### Statistical Analysis

To examine the relationships between sensorimotor behavior and non-lesioned subcortical volumes, we performed linear mixed-effects regressions. A separate regression model was run for the volume of each subcortical ROI (outcome) using sensorimotor behavior (e.g., primary sensorimotor behavior score, sensorimotor impairment, or activity limitations) as the primary predictor of interest. After ruling out collinearity (variance inflation factor ≤ 2.5), normalized age, ICV, and sex were included as fixed effects. Research cohort was included as a random effect. In chronic stroke, the effect of lesioned hemisphere was examined by including an interaction term between sensorimotor behavior and side of lesioned hemisphere to the model predicting subcortical volume. This was not examined in subacute stroke due to the smaller sample size. A likelihood ratio test (LRT) was performed to compare models with and without random effects and showed that the random effects were always significant. The regression assumptions of linearity, normality of the residuals, and homogeneity of the residual variance were checked via visual inspection of residuals versus fits plots as well as qq-plots for both individual observations and research cohorts. Potential influential values for both observations and cohorts were assessed using Cook’s distance with recommended thresholds.^30^ As we detected influential observations in almost all analyses, we re-ran the analyses using robust mixed-effect regression, which reduces the weight of influential observations in the models without excluding data.^31^ Results did not differ between original and robust regression models. The results of the robust regression models can be found in *Supplementary Materials*.

For all regression analyses, beta coefficients are presented for the predictor of interest (e.g., sensorimotor behavior, sensorimotor impairment, or activity limitations), along with the sample size (n), standard error (SE), 95% confidence interval (CI), degrees of freedom (df), standardized effect size (*d*), t-value, and uncorrected p-value. Statistical significance was adjusted for multiple comparisons across the 12 ROIs using a Bonferroni correction (p<0.004). Any significant fixed covariates are also reported.

We also compared sensorimotor behavior scores between left and right hemisphere stroke groups. The data violated the Wilkes-Shapiro test of normality for both groups (LHS: W=0.89, p<0.001, RHS: W=0.89, p<0.001). We therefore used a nonparametric Wilcoxon rank sum test to compare independent group samples.

All statistical analyses were conducted in R (version 3.6.3; R Core Team, 2020).^32^ The follow R libraries were used for the statistical analyses: the *lme* function from *nmle* was used for the linear mixed-effects regressions,^33^ the *rlmer* function from *robustlmm* was used for the robust linear mixed-effects regressions,^34^ and the *rstatix* library was used for the Wilcoxon rank sum test.^35^ In addition, *influence*.*ME* was used to detect influential values^*30*^ and *dplyr*^*36*^ and *tidyverse*^*37*^ libraries were used for data organization.

## RESULTS

Data from 828 individuals from 28 cohorts worldwide were included (see Table 1 for an overview of cohort characteristics). Briefly, the median age was 63 years old (interquartile range (IQR) 19 years), and there were 516 males and 312 females.

In subacute stroke (≤ 90 days; n=274), worse post-stroke sensorimotor behavior was significantly associated with smaller volumes of the ipsilesional thalamus (n=274, *d*=0.46, p=0.002; Table 2; Figure 1). Analysis of only individuals within just the first 21 days post-stroke (n=179, *d*=0.68, p<0.001) demonstrated the same result with a stronger effect (Table 2).

**Table 2.** Relationships between non-lesioned subcortical volumes and sensorimotor behavior in subacute and early stroke. Results from linear mixed-effects models of individuals with subacute stroke (top) and early stroke (bottom). Results in bold indicate significance with a Bonferroni correction for multiple comparisons (p<0.004). The beta coefficient for sensorimotor behavior (beta) with 95% confidence interval (CI), along with the sample size (n), standard error (SE), degrees of freedom (df), standardized effect size (*d*), t-value, and uncorrected p-value are reported, in addition to significant fixed covariates including age, sex, and intracranial volume (ICV).

| SUBACUTE STROKE ($\leq 90$ days) | | | | | | | | |
| --- | --- | --- | --- | --- | --- | --- | --- | --- |
| Brain Region | n | beta (CI) | SE | df | t-value | p-value | *d* | Significant covariates |
| *Ipsilesional* |  |  |  |  |  |  |  |  |
| Caudate | 194 | -0.01 (-0.51-0.48) | 0.25 | 180 | -0.06 | 0.954 | -0.01 | ICV |
| Lateral ventricle | 274 | 0.18 (-0.14-0.51) | 0.16 | 259 | 1.13 | 0.258 | 0.14 | Age, ICV |
| Nucleus accumbens | 245 | 0.24 (-0.14-0.62) | 0.19 | 231 | 1.26 | 0.210 | 0.17 | Age |
| Pallidum | 223 | 0.21 (-0.26-0.67) | 0.24 | 209 | 0.87 | 0.387 | 0.12 | ICV |
| Putamen | 201 | 0.39 (-0.09-0.88) | 0.25 | 187 | 1.61 | 0.109 | 0.24 | Age, ICV |
| **Thalamus** | **210** | **0.69 (0.27-1.11)** | **0.21** | **197** | **3.21** | **0.002** | **0.46** | **Age, ICV** |
| *Contralesional* |  |  |  |  |  |  |  |  |
| Caudate | 219 | 0.22 (-0.20-0.64) | 0.21 | 205 | 1.04 | 0.298 | 0.15 | ICV |
| Lateral ventricle | 274 | 0.15 (-0.18-0.49) | 0.17 | 259 | 0.92 | 0.361 | 0.11 | Age, ICV |
| Nucleus accumbens | 253 | 0.15 (-0.23-0.52) | 0.19 | 239 | 0.77 | 0.443 | 0.10 | Age, ICV |
| Pallidum | 250 | 0.50 (0.07-0.92) | 0.22 | 236 | 2.30 | 0.022 | 0.30 | ICV |
| Putamen | 229 | 0.37 (-0.05-0.79) | 0.21 | 215 | 1.75 | 0.081 | 0.24 | Age, ICV |
| Thalamus | 217 | 0.09 (-0.33-0.50) | 0.21 | 204 | 0.41 | 0.679 | 0.06 | Age, ICV |
| EARLY STROKE ($\leq 21$ days) | | | | | | | | |
| Brain Region | n | beta (CI) | SE | df | t-value | p-value | *d* | Significant covariates |
| *Ipsilesional* |  |  |  |  |  |  |  |  |
| Caudate | 135 | -0.09 (-0.67-0.48) | 0.29 | 125 | -0.32 | 0.749 | -0.06 | ICV |
| Lateral ventricle | 182 | 0.25 (-0.11-0.61) | 0.18 | 172 | 1.37 | 0.173 | 0.21 | Age, ICV |
| Nucleus accumbens | 165 | 0.19 (-0.23-0.60) | 0.21 | 155 | 0.90 | 0.369 | 0.14 | Age |
| Pallidum | 157 | 0.12 (-0.39-0.63) | 0.26 | 147 | 0.46 | 0.644 | 0.08 | ICV |
| Putamen | 143 | 0.25 (-0.28-0.79) | 0.27 | 133 | 0.93 | 0.354 | 0.16 | Age, ICV |
| **Thalamus** | **137** | **0.79 (0.38-1.20)** | **0.21** | **128** | **3.82** | **<0.001** | **0.68** | **Age, ICV** |
| *Contralesional* |  |  |  |  |  |  |  |  |
| Caudate | 147 | 0.17 (-0.29-0.64) | 0.24 | 137 | 0.74 | 0.461 | 0.13 | ICV |
| Lateral ventricle | 182 | 0.19 (-0.20-0.57) | 0.19 | 172 | 0.96 | 0.337 | 0.15 | Age, ICV |
| Nucleus accumbens | 170 | 0.30 (-0.09-0.69) | 0.20 | 160 | 1.53 | 0.127 | 0.24 | Age |
| Pallidum | 171 | 0.65 (0.19-1.11) | 0.23 | 161 | 2.79 | 0.006 | 0.44 | ICV |
| Putamen | 158 | 0.26 (-0.21-0.72) | 0.24 | 148 | 1.10 | 0.274 | 0.18 | Age, ICV |
| Thalamus | 150 | 0.20 (-0.28-0.67) | 0.24 | 141 | 0.82 | 0.411 | 0.14 | Age, ICV |

**Figure 1.**
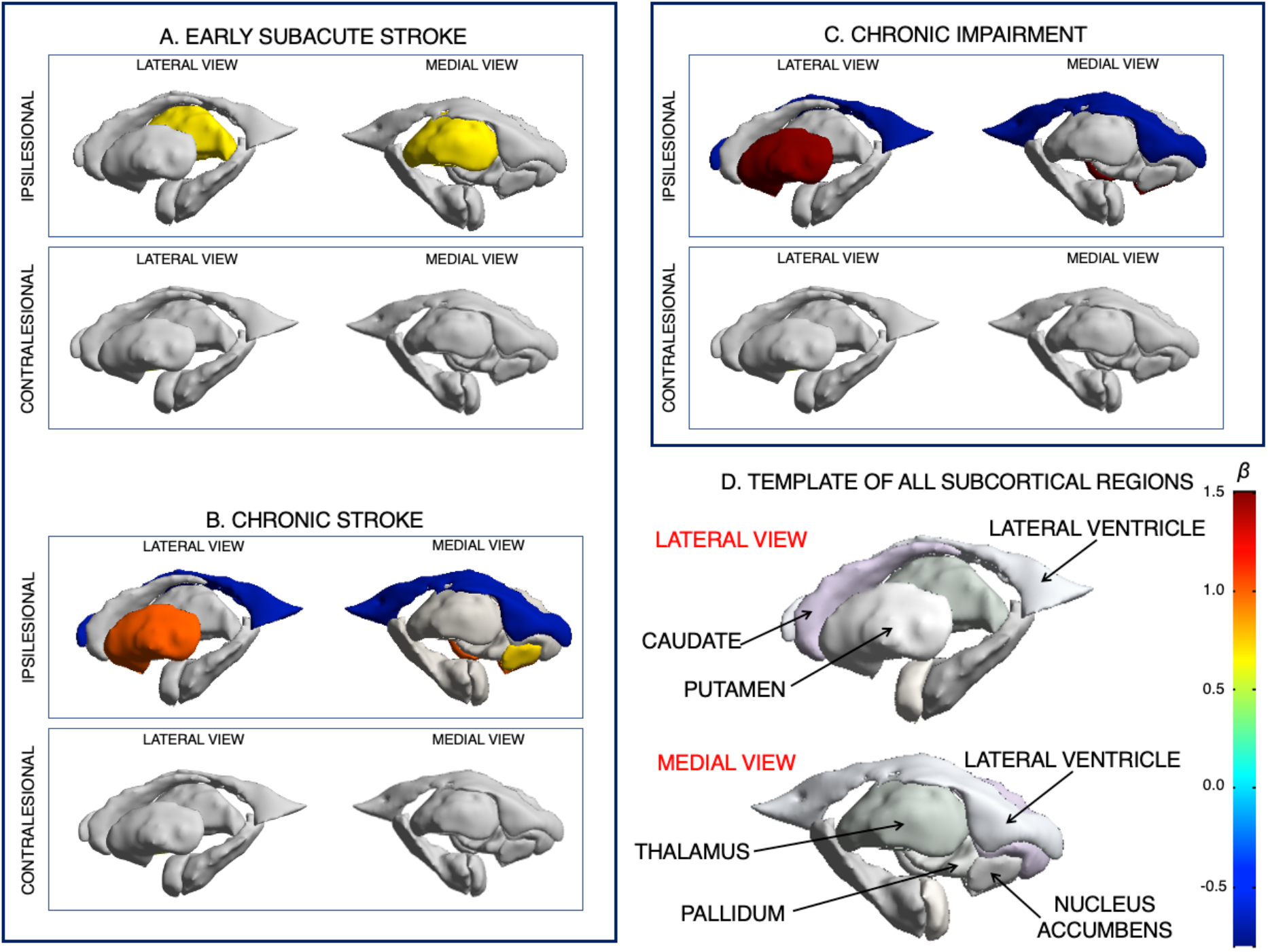
Relationships between post-stroke sensorimotor behavior and non-lesioned subcortical volumes. Non-lesioned subcortical regions (1D, bottom right) that relate to sensorimotor behavior from linear mixed-effects models of people with subacute (1A, top left) and chronic (1B, bottom left) stroke. Non-lesioned subcortical volume relationships with chronic sensorimotor impairment is shown in 1C (top right). There were no significant volume relationships with chronic activity limitations. Colors represent the beta estimate (β) for sensorimotor behavior from each model. Warmer colors represent stronger positive relationships (e.g., larger brain volumes relate to better behavior), and cooler colors represent stronger negative relationships (e.g., larger brain volumes relate to worse behavior).

In chronic stroke (≥ 180 days; n=404), worse sensorimotor behavior was related to smaller volumes of the ipsilesional putamen (*d*=0.52, p<0.001) and ipsilesional nucleus accumbens (*d*=0.39, p=0.002), and a larger volume of the ipsilesional lateral ventricle (*d*=-0.42, p<0.001; Table 3; Figure 1).

**Table 3.** Relationships between non-lesioned subcortical volumes and sensorimotor behavior in chronic stroke. Results from linear mixed-effects models of individuals with chronic stroke. Results in bold indicate significance with a Bonferroni correction for multiple comparisons (p<0.004). The beta coefficient for sensorimotor behavior (beta) with 95% confidence interval (CI), along with the sample size (n), standard error (SE), degrees of freedom (df), standardized effect size (*d*), t-value, and uncorrected p-value are reported, in addition to significant fixed covariates including age, sex, and intracranial volume (ICV).

| CHRONIC STROKE ($\geq 180$ days) | | | | | | | | |
| --- | --- | --- | --- | --- | --- | --- | --- | --- |
| Brain Region | n | beta (CI) | SE | df | t-value | p-value | d | Significant covariates |
| *Ipsilesional* |  |  |  |  |  |  |  |  |
| Caudate | 193 | 0.27 (-0.28-0.82) | 0.28 | 169 | 0.98 | 0.330 | 0.15 | ICV |
| **Lateral ventricle** | **404** | **-0.70 (-1.04--0.36)** | **0.17** | **378** | **-4.04** | **<0.001** | **-0.42** | **Age, ICV** |
| **Nucleus accumbens** | **289** | **0.72 (0.27-1.18)** | **0.23** | **264** | **3.15** | **0.002** | **0.39** | **Age** |
| Pallidum | 225 | 0.30 (-0.23-0.84) | 0.27 | 200 | 1.11 | 0.267 | 0.16 | ICV |
| **Putamen** | **207** | **1.01 (0.45-1.57)** | **0.28** | **183** | **3.54** | **<0.001** | **0.52** | **Age** |
| Thalamus | 169 | 0.08 (-0.60-0.75) | 0.34 | 146 | 0.22 | 0.827 | 0.04 | Age |
| *Contralateral* |  |  |  |  |  |  |  |  |
| Caudate | 345 | 0.08 (-0.31-0.48) | 0.20 | 320 | 0.41 | 0.679 | 0.05 | ICV |
| Lateral ventricle | 404 | -0.39 (-0.70--0.07) | 0.16 | 378 | -2.42 | 0.016 | -0.25 | Age, ICV |
| Nucleus accumbens | 344 | 0.21 (-0.22-0.65) | 0.22 | 319 | 0.96 | 0.339 | 0.11 | Age |
| Pallidum | 359 | 0.20 (-0.20-0.60) | 0.20 | 334 | 0.97 | 0.332 | 0.11 | Sex, ICV |
| Putamen | 355 | 0.21 (-0.18-0.60) | 0.20 | 330 | 1.06 | 0.291 | 0.12 | Age, ICV |
| Thalamus | 329 | -0.24 (-0.60-0.12) | 0.18 | 304 | -1.29 | 0.196 | -0.15 | Age, ICV |

In chronic stroke, we examined brain-behavior relationships using a measure of impairment (the FMA-UE scale; n=256) and two measures of activity limitation (WMFT, ARAT; n=116). Worse sensorimotor impairment was associated with smaller ipsilesional putamen (*d*=0.72, p=0.001) and larger ipsilesional lateral ventricle volumes (*d*=-0.41, p=0.002; Table 4; Figure 1). We found no significant relationships between subcortical volumes and measures of activity limitations (Table 4).

**Table 4.** Relationships between non-lesioned subcortical volumes and two measures of sensorimotor behavior (impairment, activity limitations). Results from linear mixed-effects models in individuals with chronic stroke of sensorimotor impairment (top) compared to activity limitations (bottom). Results in bold indicate significance with a Bonferroni correction for multiple comparisons (p<0.004). The beta coefficient for sensorimotor impairment/activity limitations (beta) with 95% confidence interval (CI), along with the sample size (n), standard error (SE), degrees of freedom (df), standardized effect size (*d*), t-value, and uncorrected p-value are reported, in addition to significant fixed covariates including age, sex, and intracranial volume (ICV).

| SENSORIMOTOR IMPAIRMENT IN CHRONIC STROKE |  |  |  |  |  |  |  |  |
| --- | --- | --- | --- | --- | --- | --- | --- | --- |
| Brain Region | n | beta (CI) | SE | df | t-value | p-value | d | Significant covariates |
| *Ipsilesional* |  |  |  |  |  |  |  |  |
| Caudate | 94 | 0.92 (-0.06-1.89) | 0.49 | 77 | 1.87 | 0.065 | 0.43 | ICV |
| **Lateral ventricle** | **256** | **-0.74 (-1.20--0.27)** | **0.24** | **237** | **-3.13** | **0.002** | **-0.41** | **Age, ICV** |
| Nucleus accumbens | 171 | 0.58 (0.01-1.15) | 0.29 | 153 | 2.02 | 0.045 | 0.33 | Age |
| Pallidum | 120 | 0.76 (0.01-1.51) | 0.38 | 102 | 2.02 | 0.046 | 0.40 | - |
| **Putamen** | **104** | **1.50 (0.61-2.39)** | **0.45** | **87** | **3.34** | **0.001** | **0.72** | - |
| Thalamus | 84 | 0.33 (-0.72-1.38) | 0.53 | 68 | 0.62 | 0.537 | 0.15 | - |
| *Contralesional* |  |  |  |  |  |  |  |  |
| Caudate | 222 | 0.06 (-0.44-0.57) | 0.26 | 204 | 0.25 | 0.806 | 0.03 | ICV |
| Lateral ventricle | 256 | -0.51 (-0.88--0.14) | 0.19 | 237 | -2.70 | 0.007 | -0.35 | Age, ICV |
| Nucleus accumbens | 222 | 0.21 (-0.31-0.73) | 0.26 | 204 | 0.80 | 0.425 | 0.11 | Age |
| Pallidum | 231 | 0.20 (-0.33-0.73) | 0.27 | 213 | 0.74 | 0.459 | 0.10 | Sex |
| Putamen | 229 | 0.10 (-0.38-0.58) | 0.24 | 211 | 0.41 | 0.681 | 0.06 | Age, ICV |
| Thalamus | 211 | -0.40 (-0.88-0.07) | 0.24 | 193 | -1.67 | 0.096 | -0.24 | Age, ICV |
| ACTIVITY LIMITATIONS IN CHRONIC STROKE |  |  |  |  |  |  |  |  |
| Brain Region | n | beta (CI) | SE | df | t-value | p-value | d | Significant covariates |
| *Ipsilesional* |  |  |  |  |  |  |  |  |
| Caudate | 52 | -0.63 (-1.80-0.53) | 0.58 | 44 | -1.09 | 0.280 | -0.33 | - |
| Lateral ventricle | 116 | -0.71 (-1.46-0.04) | 0.38 | 108 | -1.88 | 0.062 | -0.36 | Age, ICV |
| Nucleus accumbens | 86 | 0.77 (-0.31-1.85) | 0.54 | 78 | 1.42 | 0.159 | 0.32 | - |
| Pallidum | 64 | 0.71 (-0.25-1.67) | 0.48 | 56 | 1.47 | 0.146 | 0.39 | - |
| Putamen | 65 | 0.71 (-0.62-2.04) | 0.67 | 57 | 1.06 | 0.292 | 0.28 | - |
| Thalamus | 56 | 0.94 (-0.36-2.25) | 0.65 | 48 | 1.45 | 0.153 | 0.42 | - |
| *Contralesional* |  |  |  |  |  |  |  |  |
| Caudate | 96 | -0.07 (-0.98-0.84) | 0.46 | 88 | -0.15 | 0.885 | -0.03 | - |
| Lateral ventricle | 116 | -0.72 (-1.44-0.01) | 0.37 | 108 | -1.95 | 0.054 | -0.38 | Age, ICV |
| Nucleus accumbens | 107 | -0.34 (-1.17-0.49) | 0.42 | 99 | -0.81 | 0.420 | -0.16 | Age |
| Pallidum | 103 | -0.15 (-0.98-0.68) | 0.42 | 95 | -0.35 | 0.728 | -0.07 | Sex |
| Putamen | 100 | 0.06 (-0.91-1.03) | 0.49 | 92 | 0.12 | 0.903 | 0.03 | Age |
| Thalamus | 92 | 0.28 (-0.51-1.06) | 0.39 | 84 | 0.71 | 0.482 | 0.15 | Age, ICV |

In chronic stroke, we further analyzed the differences between individuals with left hemisphere stroke (LHS, n=214) versus right hemisphere stroke (RHS, n=190) by including lesioned hemisphere as an interaction term in the model. There were no significant effects of the side of the lesioned hemisphere on the relationship between sensorimotor behavior and subcortical volumes, and no main effects of the lesioned hemisphere (see *Supplementary Materials*). Inclusion of the lesioned hemisphere into the model did not change the main effects of sensorimotor behavior. We also examined whether there were differences in behavioral scores for LHS and RHS groups. The median sensorimotor behavior score in LHS was 0.80 (IQR=0.39) and in RHS was 0.74 (IQR=0.49). A Wilcoxon test showed no significant effect of lesioned hemisphere between groups (p=0.29, effect size r=0.053).

Finally, an exploratory analysis of the entire cohort (N=828) demonstrated significant relationships between worse sensorimotor behavior and smaller volumes of the ipsilesional thalamus (*d*=0.33, p=0.001), putamen (*d*=0.33, p<0.001), and nucleus accumbens (*d*=0.23, p=0.004), and a larger lateral ventricle volume (*d*=-0.23, p=0.001; see *Supplementary Materials*).

## DISCUSSION

We report the first international, multi-site pooled analysis with individual patient data using high-resolution structural brain imaging in stroke rehabilitation research and the largest study to date relating spared subcortical brain volumes to post-stroke sensorimotor behavior. We identified novel, significant relationships between worse post-stroke sensorimotor behavior and smaller volumes of spared deep gray matter structures, including the ipsilesional thalamus, putamen, and nucleus accumbens, as well as general atrophy as indexed by enlargement of the ipsilesional lateral ventricle. Notably, analyses included only non-lesioned structures, and significant relationships were found only in the ipsilesional hemisphere. These findings suggest that, post-stroke, secondary subcortical brain alterations related to sensorimotor behavior occur most prominently in the hemisphere directly affected by the stroke. This was observed despite the fact that, after stroke, atrophy and reorganization has been observed bilaterally.^38^ The identification of sensorimotor relationships with these specific ipsilesional subcortical nuclei may provide novel neuromodulatory or pharmacological targets to improve stroke outcomes.

Our results support the hypothesis that different non-lesioned deep gray structures serve distinct roles in subacute versus chronic stroke, which is not surprising given the cascade of neurobiological and neuroinflammatory processes that occur early after stroke.^39, 40^ Within 90 days after stroke, only the ipsilesional thalamus showed detectable associations with post-stroke sensorimotor behavior, in line with recent research showing marked thalamic atrophy, especially within the first three months post-stroke.^38^ A smaller thalamic volume could reflect cell loss and thalamic dysfunction, thereby limiting resources crucial for early recovery.^4, 38^ Importantly, we found that this relationship is not only present but stronger in the first 21 days post-stroke. As non-lesioned brain volumes within six weeks after stroke are assumed to be similar to those before the stroke,^21^ this finding suggests that larger thalamic volumes prior to stroke could provide a neuroprotective effect. Thalamic atrophy was recently associated with loss of extrinsic and intrinsic connectivity between the thalamus and the rest of the brain, suggesting that thalamic measures may serve as an index of global brain function.^41^ Future research using longitudinal datasets with greater spatial specificity could relate changes in specific thalamic nuclei to sensorimotor recovery to identify targets for neuroprotective or early stroke therapies.

In chronic stroke, reduced volumes of the ipsilesional putamen and nucleus accumbens were consistently associated with worse sensorimotor behavior. General atrophy, as indexed by a larger ipsilesional ventricle volume, was also negatively associated with sensorimotor behavioral measures. This is the first large-scale validation showing volume of these specific structures as correlates of sensorimotor behavioral outcomes in chronic stroke. This finding augments existing stroke literature, which has typically examined direct damage to combined subcortical regions, without differentiating roles of the individual basal ganglia nuclei and thalamus. Here, we specifically identify the putamen and nucleus accumbens, which are key components of corticostriatal and mesolimbic circuits, and which both represent key dopaminergic targets in the brain.

Specifically, within the corticostriatal circuit, the putamen receives direct cortical signals from the primary motor, premotor, and sensory cortices and relays them to the thalamus to modulate motor control. Interestingly, although the caudate also relays input to the thalamus, it receives its inputs from multimodal association cortices and visual regions—not primary motor regions—and did not have a significant brain-behavior relationship in our analyses. This distinction suggests that post-stroke sensorimotor behavior is primarily associated with subcortical nuclei specifically receiving direct sensorimotor input. In line with this, we found that smaller putamen volumes related to both worse sensorimotor behavior generally and impairment specifically, as evidenced by the association with the FMA-UE in chronic stroke. This finding is in line with previous work showing that direct damage to the putamen relates to post-stroke gait impairment,^42^ upper limb impairment,^43^ and spasticity,^44^ all deficits which overlap with the behavioral measures used here. In addition, secondary atrophy of the putamen has been reported after cortical stroke and is associated with infarct volume^45^ and post-stroke cognitive deficits.^46^ The relationship between chronic sensorimotor behavioral deficits and atrophy of the ipsilesional putamen after stroke, however, has not previously been reported. As atrophy of the putamen has been associated with a wide variety of neuropsychiatric and neurodegenerative disorders,^47^ including Alzheimer’s disease,^48^ multiple sclerosis, attention deficit disorder,^12^ and Huntington’s disease,^10^ it is possible that the integrity of the putamen is required not only for specifically sensorimotor behavior but also, more generally, for overall healthy brain functioning.

While the ipsilesional nucleus accumbens was significantly related to chronic sensorimotor behavior in general, it was neither related to sensorimotor impairment (FMA-UE) nor to activity limitation. However, the analyses on impairment and activity limitations had less statistical power to detect relationships. As the nucleus accumbens is a key component of the ventral striatum and implicated in dopaminergic modulation of reward-based behaviors,^49^ this region may impact more complex aspects of motor performance, such as motivation and participation, that may not be reflected in metrics of impairment or activity. A number of studies show decreases in ventral striatal processes such as reward sensitivity, motivation, and apathy after stroke,^50^ and post-stroke hypoactivity in the nucleus accumbens has been identified during reward-based decision-making tasks.^51^ Thus the nucleus accumbens may affect sensorimotor behavior by influencing reward and motivation,^52^ which could impact use of the affected limb in daily tasks. Although pharmacological methods to modulate the dopaminergic system and promote motor recovery following stroke have been widely studied, individual outcomes vary widely.^53^ Future research may investigate whether individual differences in the volume and connectivity of the nucleus accumbens predict who may benefit from dopaminergic treatment.

In chronic stroke, we also detected an association between an enlarged ipsilesional lateral ventricle and poor sensorimotor behavior. This relationship was only significant at the chronic stage and was exclusive to the ipsilesional lateral ventricle, which may be due to hydrocephalus ex vacuo. Ventricular enlargement post-stroke may also be influenced by small vessel disease (i.e., leukoaraiosis), although this is typically observed bilaterally.^19^ Enlargement of the bilateral lateral ventricles has also been associated with generalized brain atrophy that occurs during aging and with impaired cognitive function.^54^ The contrast between ipsilesional and contralesional ventricles may provide unique insight into the specific impact of the stroke versus general aging on chronic stroke sensorimotor outcomes.

Our results also suggest that there are distinct brain-behavior relationships for different ICF dimensions of sensorimotor behavior. Chronic motor impairment, as measured by the FMA-UE, was associated with a smaller ipsilesional putamen and larger ipsilesional ventricle, which may provide an indication of corticostriatal circuit integrity as well as more general brain functions essential for sensorimotor control. In contrast, there were no subcortical associations with activity limitations in the current study, possibly related to the smaller sample size. Activity limitations may also be more strongly related to the integrity or function of distributed regions across whole brain networks rather than subcortical structures,^55, 56^ given that functional performance can be influenced by psychosocial factors to a greater degree than impairment measures.

Findings did not indicate a significant effect of lesioned hemisphere on the relationship between chronic sensorimotor behavior and spared subcortical volumes. These results are surprising, given that the large majority of patients were likely left hemisphere dominant for motor control, and previous research has identified specialized hemispheric in sensorimotor control after stroke.^25^ However, previous research has primarily focused on cortical regions and functional activity, rather than subcortical structures. Side of stroke injury may not directly impact sensorimotor relationships with spared subcortical volumes.

Finally, the current results represent the first large-scale, multi-site analysis utilizing harmonized high-resolution brain imaging and behavioral measures in the field of stroke rehabilitation. The fact that the current results, using diverse stroke rehabilitation data, fit with existing literature and reveal new findings is further confirmation that such an approach is not only feasible and effective, but also beneficial for moving the stroke rehabilitation field forward.

### Limitations and Future Directions

A key limitation of pooling multi-site data is inconsistent variables across cohorts, limiting subgroup analyses and reducing the number of included covariates. Models only included the covariates age, sex, and intracranial volume; however, many additional demographic variables, such as duration and type of rehabilitation received, handedness, race, educational level, and comorbidities, may influence these relationships. In addition, larger sample sizes for different sensorimotor outcome measures would provide greater support for the current findings. Related, small high-resolution MRI samples (n < 50) at earlier time points of stroke (i.e., ≤ 7 days, defined as acute^20^) with sensorimotor behavioral outcomes limited our ability to specifically examine acute brain-behavior relationships or to examine relationships between impairment versus activity limitations in acute or subacute stroke in the current analysis. The ENIGMA Stroke Recovery Working Group recommends following consensus guidelines for greater harmonization of prospectively-collected data to facilitate more precise pooled analyses across all times after stroke.^14, 57^

Lesion overlap with subcortical regions and poor segmentation of subcortical regions due to lesion-induced distortions resulted in a variable sample size for each ROI, potentially limiting the power to detect relationships in regions with smaller samples. Furthermore, exclusion of individuals with lesioned or incorrectly segmented ROIs may have disproportionately excluded individuals with larger lesions, who may be more severely affected. This could have biased the sample towards more mild-to-moderately impaired patients. Future studies using information about the lesions (lesion location, volume, and overlap) derived from accurately segmented lesion masks for each observation could address these issues.

Finally, many of these subcortical regions are also critical for and related to post-stroke cognition, mood, sleep, learning and other traits of interest. While this analysis was limited to sensorimotor behavioral measures to maximize available data for analysis, these findings may not be unique to sensorimotor behavior. Future studies should assess the relationship between these subcortical volumes and additional stroke outcome measures.

## Conclusion

This international collaborative analysis revealed significant relationships between post-stroke sensorimotor behavior and volumetric measures of the residual ipsilesional thalamus, putamen, nucleus accumbens, and lateral ventricle at different times after stroke – brain metrics that may reflect overall brain health and network integrity and could lead to the identification of novel neural targets for pharmacological or behavioral modulation in stroke rehabilitation.

## ACKNOWLEDGMENTS

We thank all of the members of the ENIGMA Stroke Recovery working group, all of the participants, as well as Sophia Thomopolous for her assistance.

## REFERENCES

1. Krakauer JW, Carmichael ST. Broken movement: The neurobiology of motor recovery after stroke. MIT Press; 2017.

2. Hibar DP, Stein JL, Renteria ME, Arias-Vasquez A, Desrivières S, Jahanshad N, et al. Common genetic variants influence human subcortical brain structures. Nature. 2015;520:224

3. Binkofski F, Seitz R, Arnold S, Classen J, Benecke R, Freund HJ. Thalamic metabolism and corticospinal tract integrity determine motor recovery in stroke. Annals of neurology. 1996;39:460–470

4. Fries W, Danek A, Scheidtmann K, Hamburger C. Motor recovery following capsular stroke: Role of descending pathways from multiple motor areas. Brain. 1993;116:369–382

5. Shelton FtdN, Reding MJ. Effect of lesion location on upper limb motor recovery after stroke. Stroke. 2001;32:107–112

6. Kuceyeski A, Navi BB, Kamel H, Raj A, Relkin N, Toglia J, et al. Structural connectome disruption at baseline predicts 6-months post-stroke outcome. Human brain mapping. 2016;37:2587–2601

7. Jones EG. The thalamus. Springer Science & Business Media; 2012.

8. Carmichael ST, Tatsukawa K, Katsman D, Tsuyuguchi N, Kornblum HI. Evolution of diaschisis in a focal stroke model. Stroke. 2004;35:758–763

9. Alexander GE, Crutcher MD, DeLong MR. Basal ganglia-thalamocortical circuits: Parallel substrates for motor, oculomotor,”prefrontal” and “limbic” functions. Progress in brain research. Elsevier; 1991:119–146.

10. Lanciego JL, Luquin N, Obeso JA. Functional neuroanatomy of the basal ganglia. Cold Spring Harbor perspectives in medicine. 2012;2:a009621

11. Boyd LA, Winstein CJ. Providing explicit information disrupts implicit motor learning after basal ganglia stroke. Learning & memory. 2004;11:388–396

12. Campbell BC, Ma H, Ringleb PA, Parsons MW, Churilov L, Bendszus M, et al. Extending thrombolysis to 4· 5–9 h and wake-up stroke using perfusion imaging: A systematic review and meta-analysis of individual patient data. The Lancet. 2019;394:139–147

13. Goyal M, Menon BK, van Zwam WH, Dippel DW, Mitchell PJ, Demchuk AM, et al. Endovascular thrombectomy after large-vessel ischaemic stroke: A meta-analysis of individual patient data from five randomised trials. The Lancet. 2016;387:1723–1731

14. Liew S-L, Zavaliangos-Petropulu A, Jahanshad N, Lang CE, Hayward KS, Lohse K, et al. The enigma stroke recovery working group: Big data neuroimaging to study brain-behavior relationships after stroke. Human brain mapping. 2020

15. Ioannidis J. Next-generation systematic reviews: Prospective meta-analysis, individual-level data, networks and umbrella reviews. 2017

16. Berlin JA, Santanna J, Schmid CH, Szczech LA, Feldman HI. Individual patient-versus group-level data meta-regressions for the investigation of treatment effect modifiers: Ecological bias rears its ugly head. Statistics in medicine. 2002;21:371–387

17. Glasziou P, Altman DG, Bossuyt P, Boutron I, Clarke M, Julious S, et al. Reducing waste from incomplete or unusable reports of biomedical research. The Lancet. 2014;383:267–276

18. Apostolova LG, Green AE, Babakchanian S, Hwang KS, Chou Y-Y, Toga AW, et al. Hippocampal atrophy and ventricular enlargement in normal aging, mild cognitive impairment and alzheimer’s disease. Alzheimer disease and associated disorders. 2012;26:17

19. Hijdra A, Verbeeten Jr B. Leukoaraiosis and ventricular enlargement in patients with ischemic stroke. Stroke. 1991;22:447–450

20. Bernhardt J, Hayward KS, Kwakkel G, Ward NS, Wolf SL, Borschmann K, et al. Agreed definitions and a shared vision for new standards in stroke recovery research: The stroke recovery and rehabilitation roundtable taskforce. International Journal of Stroke. 2017;12:444–450

21. Egorova N, Liem F, Hachinski V, Brodtmann A. Predicted brain age after stroke. Frontiers in Aging Neuroscience. 2019;11:348

22. Azari NP, Binkofski F, Pettigrew KD, Freund HJ, Seitz RJ. Enhanced regional cerebral metabolic interactions in thalamic circuitry predicts motor recovery in hemiparetic stroke. Human brain mapping. 1996;4:240–253

23. Gauthier LV, Taub E, Mark VW, Barghi A, Uswatte G. Atrophy of spared gray matter tissue predicts poorer motor recovery and rehabilitation response in chronic stroke. Stroke. 2012;43:453–457

24. McDougall J, Wright V, Rosenbaum P. The icf model of functioning and disability: Incorporating quality of life and human development. Developmental neurorehabilitation. 2010;13:204–211

25. Sainburg RL, Maenza C, Winstein C, Good D. Motor lateralization provides a foundation for predicting and treating non-paretic arm motor deficits in stroke. Progress in motor control. Springer; 2016:257–272.

26. Fischl B, Salat DH, Busa E, Albert M, Dieterich M, Haselgrove C, et al. Whole brain segmentation: Automated labeling of neuroanatomical structures in the human brain. Neuron. 2002;33:341–355

27. Fugl-Meyer AR, Jaasko L, Leyman I, Olsson S, Steglind S. The post-stroke hemiplegic patient. 1. A method for evaluation of physical performance. Scandinavian journal of rehabilitation medicine. 1975;7:13–31

28. Lyle RC. A performance test for assessment of upper limb function in physical rehabilitation treatment and research. International journal of rehabilitation research. 1981;4:483–492

29. Wolf SL, Catlin PA, Ellis M, Archer AL, Morgan B, Piacentino A. Assessing wolf motor function test as outcome measure for research in patients after stroke. Stroke. 2001;32:1635–1639

30. Nieuwenhuis R, Te Grotenhuis H, Pelzer B. Influence. Me: Tools for detecting influential data in mixed effects models. 2012

31. Greco L, Luta G, Krzywinski M, Altman N. Analyzing outliers: Robust methods to the rescue. Nature methods. 2019;16:275–277

32. Team RC. R: A language and environment for statistical computing.. R Foundation for Statistical Computing, Vienna, Austria.. 2020

33. Pinheiro J BD, DebRoy S, Sarkar D, R Core Team. _nlme: Linear and nonlinear mixed effects models_. R package version. 2020;3:1–149

34. Koller M. Robustlmm: An r package for robust estimation of linear mixed-effects models. Journal of statistical software. 2016;75:1–24

35. Kassambara A. Rstatix: Pipe-friendly framework for basic statistical tests. 2019

36. Wickham H, Francois R, Henry L, Müller K. Dplyr: A grammar of data manipulation. R package version 1.0.2. 2020

37. Wickham H, Averick M, Bryan J, Chang W, McGowan LDA, François R, et al. Welcome to the tidyverse. Journal of Open Source Software. 2019;4:1686

38. Brodtmann A, Khlif MS, Egorova N, Veldsman M, Bird LJ, Werden E. Dynamic regional brain atrophy rates in the first year after ischemic stroke. Stroke. 2020;51:e183–e192

39. Ward NS. Restoring brain function after stroke—bridging the gap between animals and humans. Nature Reviews Neurology. 2017;13:244

40. Murphy TH, Corbett D. Plasticity during stroke recovery: From synapse to behaviour. Nature reviews neuroscience. 2009;10:861

41. Mahajan KR, Nakamura K, Cohen JA, Trapp BD, Ontaneda D. Intrinsic and extrinsic mechanisms of thalamic pathology in multiple sclerosis. Annals of Neurology. 2020

42. Alexander LD, Black SE, Patterson KK, Gao F, Danells CJ, McIlroy WE. Association between gait asymmetry and brain lesion location in stroke patients. Stroke. 2009;40:537–544

43. Lee KB, Kim JS, Hong BY, Kim YD, Hwang BY, Lim SH. The motor recovery related with brain lesion in patients with intracranial hemorrhage. Behavioural Neurology. 2015;2015

44. Cheung DK, Climans SA, Black SE, Gao F, Szilagyi GM, Mochizuki G. Lesion characteristics of individuals with upper limb spasticity after stroke. Neurorehabilitation and neural repair. 2016;30:63–70

45. Baudat C, Maréchal B, Corredor-Jerez R, Kober T, Meuli R, Hagmann P, et al. Automated mri-based volumetry of basal ganglia and thalamus at the chronic phase of cortical stroke. Neuroradiology. 2020:1–10

46. Lopes MA, Firbank MJ, Widdrington M, Blamire AM, Kalaria RN, T O’Brien J. Post-stroke dementia: The contribution of thalamus and basal ganglia changes. International psychogeriatrics. 2012;24:568

47. Luo X, Mao Q, Shi J, Wang X, Li C-SR. Putamen gray matter volumes in neuropsychiatric and neurodegenerative disorders. World journal of psychiatry and mental health research. 2019;3

48. de Jong LW, van der Hiele K, Veer IM, Houwing J, Westendorp R, Bollen E, et al. Strongly reduced volumes of putamen and thalamus in alzheimer’s disease: An mri study. Brain. 2008;131:3277–3285

49. Robbins TW, Everitt BJ. Functions of dopamine in the dorsal and ventral striatum. Seminars in Neuroscience. 1992;4:119–127

50. Rochat L, Van der Linden M, Renaud O, Epiney J-B, Michel P, Sztajzel R, et al. Poor reward sensitivity and apathy after stroke: Implication of basal ganglia. Neurology. 2013;81:1674–1680

51. Widmer M, Lutz K, Luft AR. Reduced striatal activation in response to rewarding motor performance feedback after stroke. NeuroImage: Clinical. 2019;24:102036

52. Sawada M, Kato K, Kunieda T, Mikuni N, Miyamoto S, Onoe H, et al. Function of the nucleus accumbens in motor control during recovery after spinal cord injury. Science. 2015;350:98–101

53. Gower A, Tiberi M. The intersection of central dopamine system and stroke: Potential avenues aiming at enhancement of motor recovery. Frontiers in synaptic neuroscience. 2018;10:18

54. Förstl H, Zerfaß R, Geiger-Kabisch C, Sattel H, Besthorn C, Hentschel F. Brain atrophy in normal ageing and alzheimer’s disease. The British journal of psychiatry. 1995;167:739–746

55. Van Meer MP, Van Der Marel K, Wang K, Otte WM, El Bouazati S, Roeling TA, et al. Recovery of sensorimotor function after experimental stroke correlates with restoration of resting-state interhemispheric functional connectivity. Journal of Neuroscience. 2010;30:3964–3972

56. Dong Y, Dobkin BH, Cen SY, Wu AD, Winstein CJ. Motor cortex activation during treatment may predict therapeutic gains in paretic hand function after stroke. Stroke. 2006;37:1552–1555

57. Kwakkel G, Lannin NA, Borschmann K, English C, Ali M, Churilov L, et al. Standardized measurement of sensorimotor recovery in stroke trials: Consensus-based core recommendations from the stroke recovery and rehabilitation roundtable. Neurorehabilitation and neural repair. 2017;31:784-792

