## Supplementary Materials for "Smaller spared subcortical nuclei are associated with worse post-stroke sensorimotor outcomes in 28 cohorts worldwide"

| COHORT ID | SITE | COUNTRY | N | PRIMARY SENSORIMOTOR MEASURE |
|---|---|---|---|---|
| 1 | UNIVERSITY OF SOUTHERN CALIFORNIA | USA | 39 | FMA-UE |
| 2 | UNIVERSITY OF SOUTHERN CALIFORNIA | USA | 12 | FMA-UE |
| 3 | UNIVERSITY OF SOUTHERN CALIFORNIA | USA | 14 | FMA-UE |
| 4 | UNIVERSITY OF TUEBINGEN | GERMANY | 19 | FMA-UE |
| 7 | UNIVERSITY COLLEGE LONDON | UK | 42 | Action Research Arm Test |
| 8 | UNIVERSITY OF SOUTHERN CALIFORNIA | USA | 6 | Manual Muscle Test |
| 9 | UNIVERSITY OF OSLO | NORWAY | 93 | NIHSS Motor Score |
| 10 | TIANJIN MEDICAL UNIVERSITY | CHINA | 24 | FMA-UE |
| 11 | TIANJIN MEDICAL UNIVERSITY | CHINA | 29 | FMA-UE |
| 12 | UNIVERSITY OF AUCKLAND | NEW ZEALAND | 57 | FMA-UE |
| 13 | UNIVERSITY OF AUCKLAND | NEW ZEALAND | 44 | FMA-UE |
| 15 | MEDICAL UNIVERSITY OF SOUTH CAROLINA | USA | 14 | FMA-UE |
| 17 | MEDICAL UNIVERSITY OF SOUTH CAROLINA | USA | 16 | FMA-UE |
| 18 | MEDICAL UNIVERSITY OF SOUTH CAROLINA | USA | 10 | FMA-UE |
| 19 | UNIVERSITY OF GRIEFSWALD | GERMANY | 13 | Motricity Index |
| 20 | UNIVERSITY OF GRIEFSWALD | GERMANY | 21 | Motricity Index |
| 22 | UNIVERSITY OF GRIEFSWALD | GERMANY | 17 | Bogenhausen Dysphagia Score |
| 23 | UNIVERSITY OF THE SCIENCES | USA | 13 | FMA-UE |
| 24 | EMORY UNIVERSITY | USA | 21 | Manual Muscle Test |
| 25 | UNIVERSITY OF TORONTO | CANADA | 26 | Grip Strength |
| 26 | SAO PAOLO UNIVERSITY | BRAZIL | 24 | FMA-UE |
| 28 | UNIVERSITY OF CALIFORNIA, IRVINE | USA | 26 | FMA-UE |
| 31 | UNIVERSITY OF CALIFORNIA, IRVINE | USA | 35 | FMA-UE |
| 32 | UNIVERSITY OF CALIFORNIA, IRVINE | USA | 7 | FMA-UE |
| 34 | UNIVERSITY OF SOUTHERN CALIFORNIA | USA | 15 | FMA-UE |
| 35 | HOSPITAL ISRAELITA ALBERT EINSTEIN | BRAZIL | 15 | FMA-UE |
| 38 | IRCCS SANTA LUCIA FOUNDATION | ITALY | 79 | Barthel Index |
| 41 | FLOREY INSTITUTE OF NEUROSCIENCE AND MENTAL HEALTH | AUSTRALIA | 91 | NIHSS Total |

**Supplementary Table 1. Additional cohort details.** Research sites (institutions, countries) and primary sensorimotor assessment used are listed for each cohort. FMA-UE = Fugl-Meyer Assesment of Upper Extremities.

### ROBUST REGRESSIONS FOR SUBACUTE AND CHRONIC STROKE

| SUBACUTE STROKE (≤ 90 days) | | | | | | |
|---|---|---|---|---|---|---|
| **Brain Region** | **n** | **beta (CI)** | **SE** | **t-value** | **p-value** | **Significant covariates** |
| | | | ***Ipsilesional*** | | | |
| Caudate | 194 | 0.07 (-0.37 – 0.51) | 0.22 | 0.33 | 0.742 | ICV |
| Lateral ventricle | 274 | 0.15 (-0.12 – 0.42) | 0.14 | 1.11 | 0.267 | Age, ICV |
| Nucleus accumbens | 245 | 0.18 (-0.21 – 0.57) | 0.20 | 0.90 | 0.366 | Age |
| Pallidum | 223 | 0.17 (-0.22 – 0.57) | 0.20 | 0.87 | 0.387 | ICV |
| Putamen | 201 | 0.40 (-0.06 – 0.86) | 0.23 | 1.72 | 0.086 | Age, ICV |
| **Thalamus** | **210** | **0.69 (0.26 – 1.11)** | **0.22** | **3.17** | **0.002** | **Age, ICV** |
| | | | ***Contralesional*** | | | |
| Caudate | 219 | 0.23 (-0.20 – 0.66) | 0.22 | 1.04 | 0.297 | ICV |
| Lateral ventricle | 274 | 0.06 (-0.24 – 0.36) | 0.15 | 0.37 | 0.714 | Age, ICV |
| Nucleus accumbens | 253 | 0.19 (-0.17 – 0.55) | 0.18 | 1.05 | 0.294 | Age, ICV |
| Pallidum | 250 | 0.59 (0.17 – 1.01) | 0.22 | 2.73 | 0.006 | ICV |
| Putamen | 229 | 0.31 (-0.08 – 0.70) | 0.20 | 1.57 | 0.116 | Age, ICV |
| Thalamus | 217 | 0.09 (-0.25 – 0.44) | 0.18 | 0.52 | 0.603 | Age, ICV |

| CHRONIC STROKE (≥ 180 days) | | | | | | |
|---|---|---|---|---|---|---|
| **Brain Region** | **n** | **beta (CI)** | **SE** | **t-value** | **p-value** | **Significant covariates** |
| | | | ***Ipsilesional*** | | | |
| Caudate | 193 | 0.31 (-0.21 – 0.83) | 0.26 | 1.18 | 0.240 | ICV |
| **Lateral ventricle** | **404** | **-0.71 (-1.02 – -0.41)** | **0.16** | **-4.58** | **<0.001** | **Age, ICV** |
| **Nucleus accumbens** | **289** | **0.64 (0.20 – 1.07)** | **0.22** | **2.88** | **0.004** | **Age** |
| Pallidum | 225 | 0.50 (0.04 – 0.97) | 0.24 | 2.13 | 0.033 | ICV |
| **Putamen** | **207** | **1.02 (0.48 – 1.55)** | **0.27** | **3.72** | **<0.001** | **Age, ICV** |
| Thalamus | 169 | 0.11 (-0.53 – 0.75) | 0.33 | 0.33 | 0.740 | Age |
| | | | ***Contralesional*** | | | |
| Caudate | 345 | 0.14 (-0.27 – 0.55) | 0.21 | 0.67 | 0.501 | ICV |
| Lateral ventricle | 404 | -0.36 (-0.63 – -0.09) | 0.14 | -2.62 | 0.009 | Age, ICV |
| Nucleus accumbens | 344 | 0.22 (-0.19 – 0.63) | 0.21 | 1.07 | 0.285 | Age |
| Pallidum | 359 | 0.32 (-0.01 – 0.64) | 0.17 | 1.91 | 0.056 | Age, ICV |
| Putamen | 355 | 0.22 (-0.15 – 0.60) | 0.19 | 1.16 | 0.244 | Age, ICV |
| Thalamus | 329 | -0.20 (-0.57 – 0.17) | 0.19 | -1.06 | 0.288 | Age, ICV |

**Supplementary Table 2. Robust regressions to examine relationships between non-lesioned subcortical volumes and sensorimotor behavior in subacute and chronic stroke.** Results from robust linear mixed-effects models of individuals with subacute stroke (top) and chronic stroke (bottom). Results in bold indicate significance with a Bonferroni correction for multiple comparisons ($p<0.004$). The beta coefficient for sensorimotor behavior (beta) with 95% confidence interval (CI), along with the sample size (n), standard error (SE), t-value, and uncorrected p-value are reported, in addition to significant fixed covariates including age, sex, and intracranial volume (ICV).

### ROBUST REGRESSIONS FOR
### CHRONIC SENSORIMOTOR IMPAIRMENT AND ACTIVITY LIMITATIONS

**CHRONIC SENSORIMOTOR IMPAIRMENT**

| **Brain Region** | **n** | **beta (CI)** | **SE** | **t-value** | **p-value** | **Significant covariates** |
|---|---|---|---|---|---|---|
| | | | ***Ipsilesional*** | | | |
| Caudate | 194 | 0.98 (0.08 – 1.88) | 0.46 | 2.14 | 0.032 | ICV |
| **Lateral ventricle** | **274** | **-0.70 (-1.13 – -0.27)** | **0.22** | **-3.2** | **0.001** | **Age, ICV** |
| Nucleus accumbens | 245 | 0.63 (0.08 – 1.18) | 0.28 | 2.24 | 0.025 | Age |
| Pallidum | 223 | 0.79 (0.07 – 1.51) | 0.37 | 2.16 | 0.031 | - |
| **Putamen** | **201** | **1.54 (0.60 – 2.47)** | **0.48** | **3.23** | **0.001** | **-** |
| Thalamus | 210 | 0.56 (-0.33 – 1.44) | 0.45 | 1.23 | 0.22 | - |
| | | | ***Contralesional*** | | | |
| Caudate | 219 | 0.02 (-0.52 – 0.56) | 0.27 | 0.08 | 0.939 | ICV |
| Lateral ventricle | 274 | -0.46 (-0.80 – -0.13) | 0.17 | -2.72 | 0.006 | Age, ICV |
| Nucleus accumbens | 253 | 0.22 (-0.32 – 0.76) | 0.28 | 0.80 | 0.423 | Age |
| Pallidum | 250 | 0.31 (-0.13 – 0.75) | 0.22 | 1.39 | 0.166 | ICV |
| Putamen | 229 | 0.02 (-0.46 – 0.51) | 0.25 | 0.09 | 0.932 | Age, ICV |
| Thalamus | 217 | -0.34 (-0.84 – 0.15) | 0.25 | -1.36 | 0.173 | Age, ICV |

**ACTIVITY LIMITATIONS**

| **Brain Region** | **n** | **beta (CI)** | **SE** | **t-value** | **p-value** | **Significant covariates** |
|---|---|---|---|---|---|---|
| | | | ***Ipsilesional*** | | | |
| Caudate | 193 | -0.48 (-1.64 – 0.69) | 0.60 | -0.80 | 0.425 | - |
| Lateral ventricle | 404 | -0.70 (-1.32 – -0.09) | 0.31 | -2.24 | 0.025 | Age, ICV |
| Nucleus accumbens | 289 | 0.68 (-0.30 – 1.67) | 0.50 | 1.36 | 0.172 | - |
| Pallidum | 225 | 0.88 (-0.04 – 1.80) | 0.47 | 1.88 | 0.060 | - |
| Putamen | 207 | 0.87 (-0.32 – 2.05) | 0.60 | 1.43 | 0.152 | - |
| Thalamus | 169 | 1.19 (-0.08 – 2.46) | 0.65 | 1.84 | 0.066 | - |
| | | | ***Contralesional*** | | | |
| Caudate | 345 | 0.14 (-0.74 – 1.02) | 0.45 | 0.32 | 0.750 | ICV |
| Lateral ventricle | 404 | -0.72 (-1.31 – -0.13) | 0.30 | -2.38 | 0.017 | Age, ICV |
| Nucleus accumbens | 344 | -0.33 (-1.13 – 0.46) | 0.41 | -0.82 | 0.413 | Age |
| Pallidum | 359 | -0.07 (-0.93 – 0.79) | 0.44 | -0.16 | 0.874 | Sex |
| Putamen | 355 | 0.33 (-0.54 – 1.20) | 0.44 | 0.75 | 0.454 | Age |
| Thalamus | 329 | 0.19 (-0.57 – 0.95) | 0.39 | 0.49 | 0.626 | Age, Sex |

**Supplementary Table 3. Robust regressions to examine relationships between non-lesioned subcortical volumes and two measures of sensorimotor behavior (impairment, activity limitations).** Results from robust linear mixed-effects models in individuals with chronic stroke showing sensorimotor impairment (top) compared to activity limitations (bottom). Results in bold indicate significance with a Bonferroni correction for multiple comparisons ($p<0.004$). The beta coefficient for sensorimotor behavior (beta) with 95% confidence interval (CI), along with the sample size (n), standard error (SE), t-value, and uncorrected p-value are reported, in addition to significant fixed covariates including age, sex, and intracranial volume (ICV).

| Brain Region | n | df | Interaction between Sensorimotor Behavior & Lesioned Hemisphere | | | Main Effect Lesioned Hemisphere | | | Main Effect Sensorimotor Behavior | | | | Significant Covariates |
|---|---|---|---|---|---|---|---|---|---|---|---|---|---|
| | | | beta | SE | p-value | beta | SE | p-value | beta | SE | p-value | *d* | |
| *Ipsilesional* | | | | | | | | | | | | | |
| Caudate | 193 | 167 | 0.14 | 0.52 | 0.789 | -0.17 | 0.44 | 0.693 | 0.26 | 0.28 | 0.365 | 0.14 | ICV |
| **Lateral ventricle** | 404 | 376 | 0.21 | 0.29 | 0.487 | -0.13 | 0.23 | 0.572 | **-0.70** | **0.17** | **<0.001** | **-0.42** | Age, ICV |
| **Nucleus accumbens** | 289 | 262 | 0.21 | 0.39 | 0.593 | -0.13 | 0.31 | 0.684 | **0.73** | **0.23** | **0.002** | **0.39** | Age |
| Pallidum | 225 | 198 | -0.53 | 0.48 | 0.272 | 0.52 | 0.39 | 0.191 | 0.35 | 0.27 | 0.207 | 0.18 | ICV |
| **Putamen** | 207 | 181 | -0.03 | 0.53 | 0.953 | -0.28 | 0.44 | 0.525 | **0.90** | **0.29** | **0.002** | **0.47** | ICV |
| Thalamus | 169 | 144 | 0.17 | 0.63 | 0.792 | -0.95 | 0.51 | 0.065 | 0.05 | 0.32 | 0.887 | 0.02 | Age, ICV |
| *Contralesional* | | | | | | | | | | | | | |
| Caudate | 345 | 318 | 0.12 | 0.33 | 0.731 | -0.14 | 0.26 | 0.583 | 0.06 | 0.21 | 0.760 | 0.03 | ICV |
| Lateral ventricle | 404 | 376 | 0.27 | 0.28 | 0.343 | 0.06 | 0.21 | 0.789 | -0.35 | 0.16 | 0.030 | -0.22 | Age, ICV |
| Nucleus accumbens | 344 | 317 | 0.38 | 0.35 | 0.282 | -0.32 | 0.27 | 0.236 | 0.18 | 0.22 | 0.423 | 0.09 | Age |
| Pallidum | 359 | 332 | 0.07 | 0.32 | 0.819 | -0.43 | 0.25 | 0.083 | 0.09 | 0.20 | 0.672 | 0.05 | Sex, ICV |
| Putamen | 355 | 328 | -0.08 | 0.31 | 0.796 | 0.14 | 0.24 | 0.553 | 0.23 | 0.20 | 0.243 | 0.13 | Age, ICV |
| Thalamus | 329 | 302 | 0.25 | 0.30 | 0.405 | 0.47 | 0.23 | 0.045 | -0.17 | 0.18 | 0.353 | -0.11 | Age, ICV |

**Supplementary Table 4. Relationships between non-lesioned subcortical volumes, lesioned hemisphere, and sensorimotor behavior in chronic stroke.** Results from linear mixed-effects models including an interaction term between lesioned hemisphere and sensorimotor behavior in people with chronic stroke. There were no significant interactions or main effects of lesioned hemisphere. The significant main effects for sensorimotor behavior remained, similar to those shown in Table 2. Results in bold indicate significance with a Bonferroni correction for multiple comparisons ($p<0.004$). The beta coefficients (beta), standard error (SE), and uncorrected p-value for the interaction between lesioned hemisphere and sensorimotor behavior, as well as main effects for lesioned hemisphere and sensorimotor behavior, are reported, along with the sample size (n), degrees of freedom (df), standardized effect size (d), and significant fixed covariates including age, sex, and intracranial volume (ICV).

### ALL STROKE

| Brain Region | n | beta (CI) | SE | df | t-value | p-value | *d* | Significant covariates |
|---|---|---|---|---|---|---|---|---|
| | | | | ***Ipsilesional*** | | | | |
| Caudate | 482 | 0.15 (-0.18-0.48) | 0.17 | 451 | 0.89 | 0.375 | 0.08 | Sex, ICV |
| **Lateral ventricle** | **828** | **-0.39 (-0.63--0.16)** | **0.12** | **796** | **-3.26** | **0.001** | **-0.23** | **Age, ICV** |
| **Nucleus accumbens** | **655** | **0.41 (0.13-0.68)** | **0.14** | **624** | **2.91** | **0.004** | **0.23** | **Age** |
| Pallidum | 546 | 0.29 (-0.04-0.61) | 0.16 | 515 | 1.75 | 0.081 | 0.15 | ICV |
| **Putamen** | **490** | **0.64 (0.28-1.00)** | **0.18** | **459** | **3.53** | **<0.001** | **0.33** | **Age, ICV** |
| **Thalamus** | **462** | **0.58 (0.25-0.91)** | **0.17** | **433** | **3.47** | **0.001** | **0.33** | **Age, ICV** |
| | | | | ***Contralesional*** | | | | |
| Caudate | 689 | 0.01 (-0.26-0.28) | 0.14 | 658 | 0.08 | 0.939 | 0.01 | ICV |
| Lateral ventricle | 828 | -0.26 (-0.49--0.03) | 0.11 | 796 | -2.27 | 0.024 | -0.16 | Age, ICV |
| Nucleus accumbens | 727 | 0.10 (-0.15-0.36) | 0.13 | 696 | 0.78 | 0.436 | 0.06 | Age, ICV |
| Pallidum | 743 | 0.28 (0.02-0.54) | 0.13 | 712 | 2.08 | 0.038 | 0.16 | Age, ICV |
| Putamen | 704 | 0.19 (-0.07-0.45) | 0.13 | 673 | 1.45 | 0.147 | 0.11 | Age, ICV |
| Thalamus | 663 | -0.02 (-0.28-0.24) | 0.13 | 633 | -0.13 | 0.898 | -0.01 | Age, ICV |

**Supplementary Table 5. Relationships between non-lesioned subcortical volumes and sensorimotor behavior post-stroke across all times after stroke (N=828).** Results from linear mixed-effects models of individuals across all times after stroke. Uncorrected p-values shown. Results in bold indicate significance with a Bonferroni correction for multiple comparisons ($p<0.004$). The beta coefficient for sensorimotor behavior (beta) with 95% confidence interval (CI), along with the sample size (n), standard error (SE), degrees of freedom (df), standardized effect size (*d*), t-value, and uncorrected p-value are reported, in addition to significant fixed covariates including age, sex, and intracranial volume (ICV).
